# Psilocybin rescues sociability deficits in an animal model of autism

**DOI:** 10.1101/2020.09.09.289348

**Authors:** Irene Mollinedo-Gajate, Chenchen Song, Marcos Sintes-Rodriguez, Tobias Whelan, Anaïs Soula, Aslihan Selimbeyoglu, Shaun Hurley, Thomas Knöpfel

**Affiliations:** Laboratory for Neuronal Circuit Dynamics, Department of Brain Sciences, Imperial College London, London, UK; COMPASS Pathways Ltd., London, UK

## Abstract

Autism spectrum disorder (ASD) is characterized by core deficits in social interaction. The classic serotonergic psychedelic psilocybin has been suggested as a therapeutic agent that may ameliorate in the core symptomology of ASD. We found that the acute response to psilocybin was attenuated in the prenatal valproic acid exposure mouse model of ASD, and importantly, psilocybin rescued the social behavioural abnormalities present in these ASD model mice.

## Main

Autism spectrum disorder (ASD) is a heterogenous neurodevelopmental disorder characterized by core deficits in social interaction and communication, repetitive or restricted behaviour and interests, and altered sensory sensitivity^1^. The aetiology and pathophysiology of ASD remains largely unknown, and is considered to be multi-factorial, encompassing both genetic and environmental factors^2-4^. The lack of a single molecular mechanism underlying the aetiology of ASD has contributed to the challenge of finding efficacious therapeutic interventions.

The regulation and functionality of the serotonergic (5-HT) system has been implicated to play an important role in the core pathophysiology of ASD^5-9^, hence its modulation is an evidence-based therapeutic target^10-12^. Psilocybin is produced by several species of mushrooms, and is one of the compounds responsible for the psychedelic effects that these mushrooms induce, likely via altering 5-HT signalling through 5-HT_2A_ receptors^13,14^. Reduced cortical 5-HT_2A_ receptor expression has been reported in ASD suggesting that this population may be less sensitive to the subjective effects of psilocybin^8,9^. Psychedelics have recently shown potential safety and efficacy for alleviating various neuropsychiatric disorders^15^. Recent human studies also showed that psilocybin can produce sustained prosocial behaviour^16,17^, and affects several facets of social cognition relevant to deficits in ASD^18,19^.

Preclinical evaluation of psilocybin’s potential to ameliorate altered sociability requires a suitable animal model that recapitulates aspects of the social deficits observed in ASD. One such model, based on prenatal exposure to valproic acid (VPA), is particularly attractive as it does not bias towards a single genetic alteration as in genetic models of ASD and thus may better represent idiopathic ASD. VPA is clinically used for the treatment of epileptic seizures, bipolar mania, and as a migraine prophylactic. Through this long-standing clinical use, it has been observed that pregnant mothers treated with VPA are at an increased risk of giving birth to children that are diagnosed with ASD^20^. In mice, prenatal VPA exposure also results in offspring exhibiting many ASD-like behavioural phenotypes^21-24^, including altered social behaviour^25-27^. Here we explored the potential of psilocybin to ameliorate altered sociability in the prenatal VPA mouse model of autism. Offspring that have been exposed to VPA *in utero* are hereon referred to as VPA group, and those exposed to saline *in utero* as Control group. VPA group displayed a modestly reduced body weight (**Extended Data Fig. 1**).

**Fig. 1:**
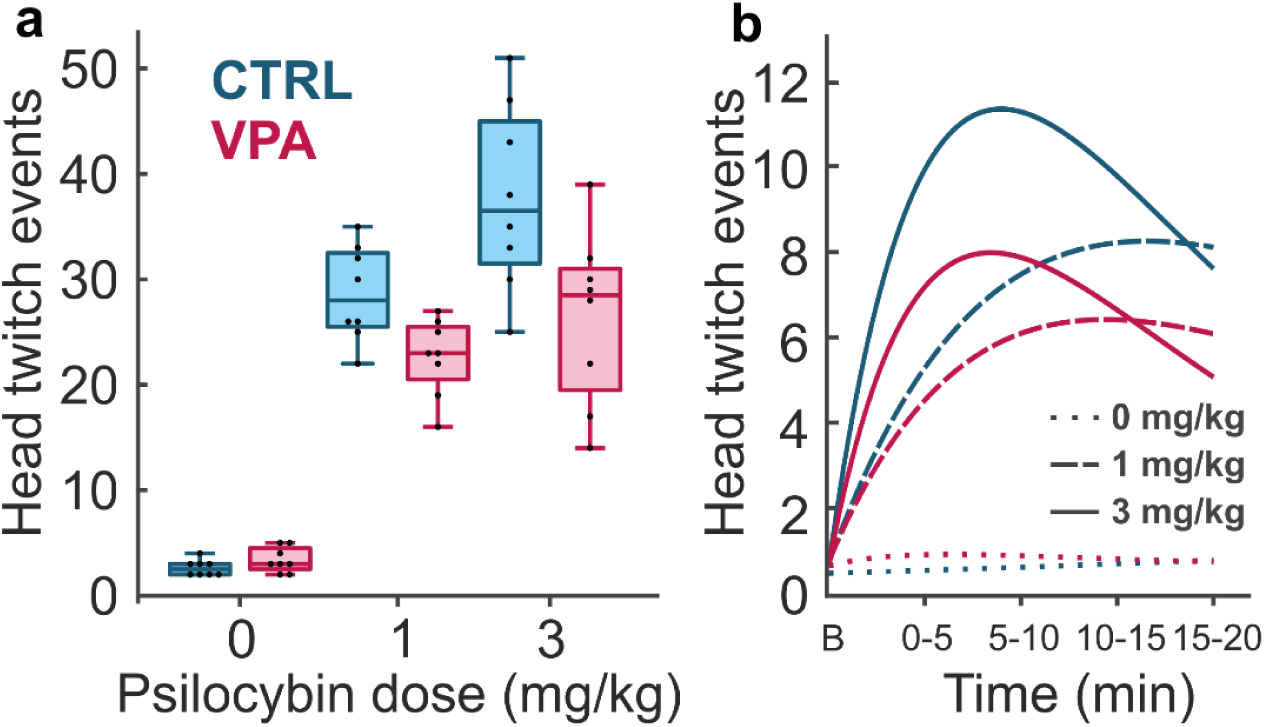
VPA group shows reduced head twitch response induced by psilocybin. **a**, Total count of head twitch events over 20 min, induced by an intraperitoneal injection of saline or psilocybin in Control and VPA groups. Psilocybin evoked dose-dependent HTR in both groups. HTR count was significantly lower with no significant dose-dependency in VPA mice (*F*_pre-treatment x treatment(2,42)_ = 4.895, *P* = 0.012). *Boxplot: center line, median; box limits, upper and lower quartiles; whiskers, 1*.*5x interquartile range*. **b**, Fitted time course of head twitch events above baseline (B) in 5-min time bins. VPA and Control groups showed similar HTR time course shapes. n = 8/group. For detail see **Extended Data Fig. 2, Supplementary Table. 1**.

We first examined the acute effect of psilocybin on both VPA and Control groups by measuring the head twitch response (HTR), a 5-HT_2A_ receptor-mediated measure that predicts hallucinogenic potency in humans^28^. All test mice (n = 8 / group) received a single intraperitoneal injection of psilocybin (low dose at 1 mg/kg or high dose at 3 mg/kg) or saline, and we counted head twitch events immediately after injections. Psilocybin evoked HTRs both in VPA and Control groups with dose dependency (**Fig. 1**). The efficacy of psilocybin to trigger HTR was significantly lower in VPA group than Control group (F_pre-treatment x treatment(2,42)_ = 4.895, *P* = 0.012, two-way ANOVA; **Fig. 1a**). The time course of the effects induced by psilocybin on both VPA and Control groups showed similar rise and decay times (**Fig. 1b, Extended Data Fig. 2**). There were no gender differences in VPA and Control groups treated with psilocybin or saline (*F*_pre-treatment x treatment x gender(2,36)_ = 0.03, *P* = 0.969, three-way ANOVA).

**Fig. 2:**
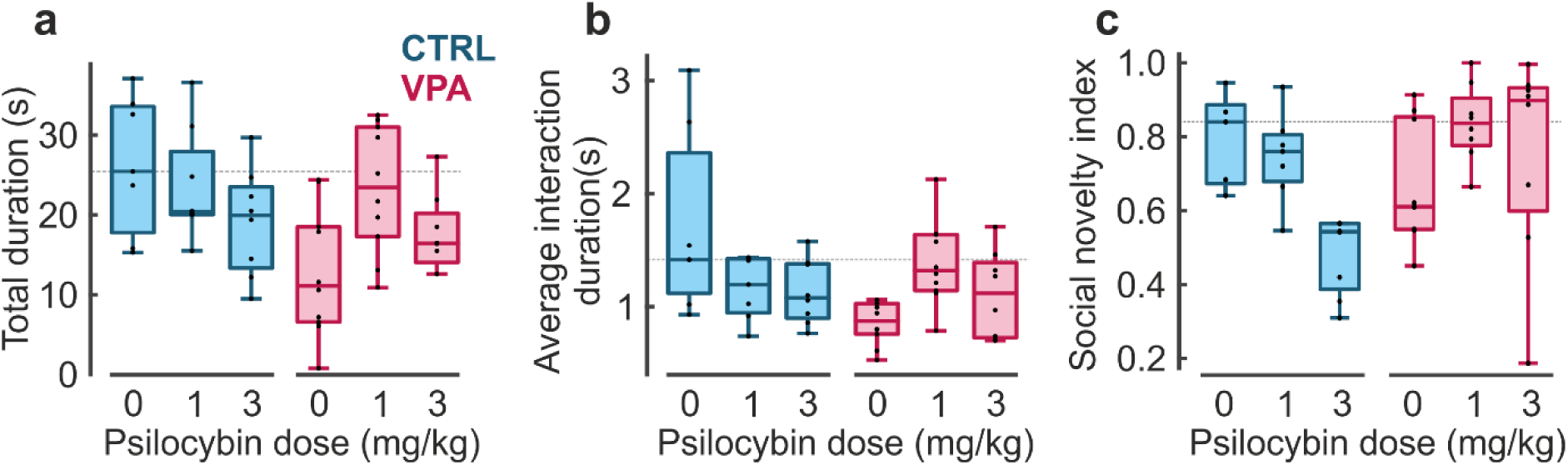
Single administration of psilocybin improves sociability and social memory in ASD model mice. **a**, Total duration time of direct nose to nose interactions during 10-min observation period (*F*_pre-treatment x treatment(2,45)_ = 3.633, *P* = 0.034), 24 hours following a single administration of saline or psilocybin in Control and VPA groups. **b**, Average time of direct nose to nose interactions (*F*_pre-treatment x treatment(2,44)_ = 6.657, *P* = 0.003), related to observations in **a. c**, Social novelty index is dose-dependently reduced in the Control group 24 hours following a single administration of saline or psilocybin (*F*_pre-treatment x treatment(2,37)_ = 6.077, *P*= 0.0052), but is increased in the VPA group. Two-way ANOVA: *n* = 7 control saline, *n* = 7 control PSI 1, *n* = 8 control PSI 3, *n* = 10 VPA saline, *n* = 10 VPA PSI 1, *n* = 8 VPA PSI 3. Dotted line represents median value from the Control group administered with saline. *Boxplot: center line, median; box limits, upper and lower quartiles; whiskers, 1*.*5x interquartile range; outliers not shown*. For statistics see **Supplementary Table. 1**

24-hour after psilocybin/saline treatments, animals were tested for their social behaviour (sociability and social memory), to capture effects beyond the acute action of psilocybin as well as to avoid acute psychedelic-like effects confounding social behaviour. VPA group spent significantly less time engaging in social interaction compared to Control group, as quantified by the total duration of nose-to-nose interactions with the stranger mouse (VPA_SAL_ vs CTRL_SAL_: *t* = 3.282, *P* = 0.005; **Fig. 2a**). A low dose of psilocybin (1 mg/kg i.p.) restored the sociability behaviour of VPA group to a level similar to that of Control group (VPA_SAL_ vs VPA_PSI1_: *t* = 2.928, *P* = 0.009; **Fig. 2a**), but the rescue effect of psilocybin was not evident following the higher dose (3 mg/kg; VPA_SAL_ vs VPA_PSI3_: *t* = 1.071, *P* = 0.300; **Fig. 2a**). Surprisingly, psilocybin treatment in the Control group showed a dose-dependent trend in reducing the total nose-to-nose interaction duration, although this effect was not statistically significant (CTRL_SAL_ vs CTRL_PSI1_: *t* = 0.657, *P* = 0.523; CTRL_SAL_ vs CTRL_PSI3_: *t* = 1.799, *P* = 0.953; **Fig. 2a**). The reduced total nose-to-nose interaction duration in VPA group was due to reduced average duration of individual interactions (VPA_SAL_ vs CTRL_SAL_: *t* = 2.952, *P* = 0.010). 1 mg/kg psilocybin substantially restored this component of social interaction behaviour and increased the duration of individual interactions when compared with the VPA saline group (VPA_SAL_ vs VPA_PSI1_: *t* = 3.307, *P* = 0.004; **Fig. 2b**). However, we did not observe any difference in the average time per interaction between VPA mice treated with 3 mg/kg psilocybin versus saline controls (VPA_SAL_ vs VPA_PSI3_: *t* = 1.326, *P* = 0.203). The surprising trend (though statistically non-significant) of psilocybin-induced reduction in social interaction in Control group was also observed in a dose-dependent trend in the average duration of individual interactions in both high and low dose psilocybin treated control mice when compared to the control saline group (CTRL_SAL_ vs CTRL_PSI1_: *t* = 1.698, *P* = 0.115; CTRL_SAL_ vs CTRL_PSI3_: *t* = 1.911, *P* = 0.078).

Next, we assessed the effect of psilocybin on social novelty preference by introducing a new stranger mouse (S2) in addition to the presence of the existing stranger mouse (S1). The social novelty index derived from this measure quantifies the extent to which S2 is recognized as new. As this recognition required remembering S1, this index is also a measure of short-term social memory. The social novelty index was reduced for the VPA group, and a single treatment of psilocybin (at either low or high doses) in VPA group increased their social novelty preference to a level similar (higher) than that of the Control group (**Fig. 2c, Supplementary Table. 1**). The opposite effect was observed in the Control group treated with either 1 mg/kg or 3 mg/kg psilocybin, where psilocybin tended to decrease the social novelty index (**Fig. 2c, Supplementary Table. 1**).

Psychedelic research is currently undergoing a renaissance, with the therapeutic potential of these compounds being increasingly recognised. Psilocybin, for example, has recently received “breakthrough designation” by the FDA in treatment-resistant depression^29^ and major depressive disorder^30^. Towards a reverse translational approach, here we report the efficacy of psilocybin in an animal model of ASD as evidence that psilocybin may be a promising pharmacotherapy for altered sociability in ASD. More detailed mechanistic studies will be needed in future to understand the molecular and circuit underpinnings of this therapeutic potential.

## Methods

### Animals

All experimental procedures were performed at Imperial College London UK in accordance with the United Kingdom Animal Scientific Procedures Act (1986), under Home Office Personal and Project Licences following appropriate ethical review. Time-mated C57BL/6J mice (Charles River, UK) were housed in groups, and separated two days prior to the expected littering date. Offspring were weaned at P28 and housed in groups of up to five animals per cage after weaning. All animals were maintained in ventilated cages, on a 12/12 h light/dark cycle at 21 ± 2°C and 55 ± 10% humidity. Water and food were provided *ad libitum*. Eight weeks old mice of both genders were used for behavioural experiments. All experiments were conducted during the daytime.

### Prenatal VPA treatment

Valproic acid (2-propylpentanoic acid) sodium salt was obtained from Sigma Aldrich (P4543; Germany), and freshly dissolved in sterile saline (0.9% NaCl) to 50 mg/ml, pH 7.4. On embryonic day 10.5 (E10.5), pregnant females were intraperitoneally administered with either a single dose of 500 mg/kg VPA or equal volume of saline. The dose and the embryonic day of VPA administration were based on previously published reports^21-23^. Prenatally exposed VPA animals showed significantly lower body weight than the prenatal saline control at weaning age (controls = 15.01 g, VPA = 12.09 g; *P* = 0.0002; n = 24/group; **Extended Data Fig. 1, Supplementary Table. 1**) and remained lower at week 8 (controls = 20.95 g, VPA = 18.40 g; *P* = 0.009; n = 24/group; **Extended Data Fig. 1, Supplementary Table. 1**). Additionally, physical abnormalities were observed in the prenatal VPA animals: 60% showed kinked tail, 10% had moderate facial abnormalities, 6% displayed fur problems, 3% had misaligned teeth. These abnormalities were not observed in the control group.

### Drug administration

Psilocybin ([3-(2-dimethylaminoethyl)-1*H*-indol-4-yl] dihydrogen phosphate, Onyx Scientific Ltd, UK) was dissolved in sterile saline solution (0.9% NaCl) to appropriate concentrations.

For behavioural experiments, 8-week old offspring were intraperitoneally injected either with saline or psilocybin (1 mg/kg, PSI 1; or 3 mg/kg, PSI 3) 24 hours prior to the sociability and social memory test. The treatments led to six different test groups: control saline, control psilocybin 1 mg/kg, control psilocybin 3 mg/kg, VPA saline, VPA psilocybin 1 mg/kg, VPA psilocybin 3 mg/kg.

### Head twitch response

Head twitch events were evaluated immediately after the i.p. administration of either saline or psilocybin for a period of 20 minutes. The mouse was placed into a customized behavioural box immediately after injection and was free to explore. The number of head-twitch events were counted by direct observation in 5-minute bins. The box (25.5 × 12.5 × 12.5 cm [L x W x H]) was made of red Plexiglass and the floor was covered with clean sawdust.

### Sociability and social memory behavioural test

Sociability and social novelty preference experiments were performed in a custom-built transparent red Plexiglass box with matt white acrylic floor. The total arena (60 × 40 × 22 cm [L × W × H]) was divided in three smaller evenly sized chambers (20 × 40 × 22 cm [L × W × H]) interconnected by 4 × 4 cm doors cut out of both central walls. Clear Plexiglass cylinders (10.5 cm internal diameter, 11 cm external diameter, 16 cm length) with 1-cm slits and 0.5-cm rods were used as cups. A red acrylic lid was placed on top of both cups to prevent animals from escaping or falling. The box was placed in a dark and quite room, illuminated from above with infrared LEDs located 1 meter over the arena.

Each behavioural testing experiment took place over 4 consecutive days (**Extended Data Fig. 1**). Animals were acclimatised to the testing room for at least one hour before the start of experiments on each day. Each test mouse was placed into the central chamber and allowed to freely explore the three-chamber setup for 10 minutes over 3 consecutive days. During this pre-habituation time, doors to the side chambers were kept open and empty cups were placed into the two side chambers at similar positions as the final test day (day 4). Cup mice were individually pre-habituated to the cups also for 10 minutes over 3 consecutive days. On day 4, the three-chamber test was performed in three phases (Habituation, Sociability, Social Memory). Each test animal was allowed to freely explore all three chambers for 10 minutes with empty cups (Habituation Phase), then driven to the central chamber with the doors closed while an age-, size-, and gender-matched unfamiliar mouse (stranger 1) was being placed into one of the cups. The test mouse was then allowed to freely explore chambers for 10 minutes (Sociability Phase). The test mouse was then returned to the central chamber while a second unfamiliar mouse (stranger 2) was placed in the remaining empty cup. Finally, the test mouse was allowed to move freely across the three chambers for 10 minutes (Social Memory Phase).

All behavioural sessions were video-recorded using a CMOS camera (Basler acA2000-165umNIR) and Basler software (Basler AG, Germany). Video recordings were used to track the position of the body of the animal in each chamber, as well as to score the duration time of direct nose to nose interactions and interaction events by a blinded well-trained observer.

### Data analysis and statistics

No statistical methods were used to predetermine sample size, but our sample size was selected based on standards in the field. Experiments were blinded to VPA or control prenatal treatment, as well as treatment during automatically or manually analysis. If applicable, outliers were identified and excluded using a Grubb’s test. Differences between two groups were compared using unpaired two-tailed Student’s *t*-test when data were normally distributed, otherwise the Mann-Whitney U test was used. Multiple group differences were assessed using two- and three-way analysis of variance (ANOVA). Independent variables were defined as: pre-treatment (Control or VPA), treatment (PSI1: psilocybin 1 mg/kg, or PSI3: psilocybin 3 mg/kg, or SAL: saline), gender. Multiple comparisons were further assessed by Bonferroni’s post hoc test. Definition of statistical significance was set at an alpha value of 0.05. Specific *P* values are reported for each analysis in the corresponding figure legend and Supplementary Table 1. All statistical analyses were performed using GraphPad Prism 8 (San Diego, CA, USA) and InVivoStat (Cambridge, United Kingdom).

Social novelty index was calculated as the following:

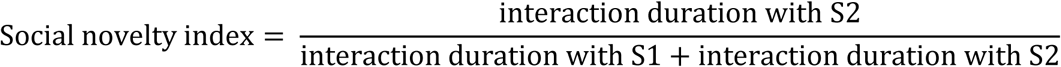

HTR responses were fitted using MATLAB (R2019b) with a single dose pharmacokinetics curve with the following equation:

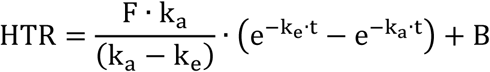

Where: HTR = head twitch response; F = drug factor (dose and bioavailability); k_a_ = absorption rate constant, k_e_ = elimination rate constant; t = time; B = baseline count.

## Extended data

**Extended Data Fig. 1:**
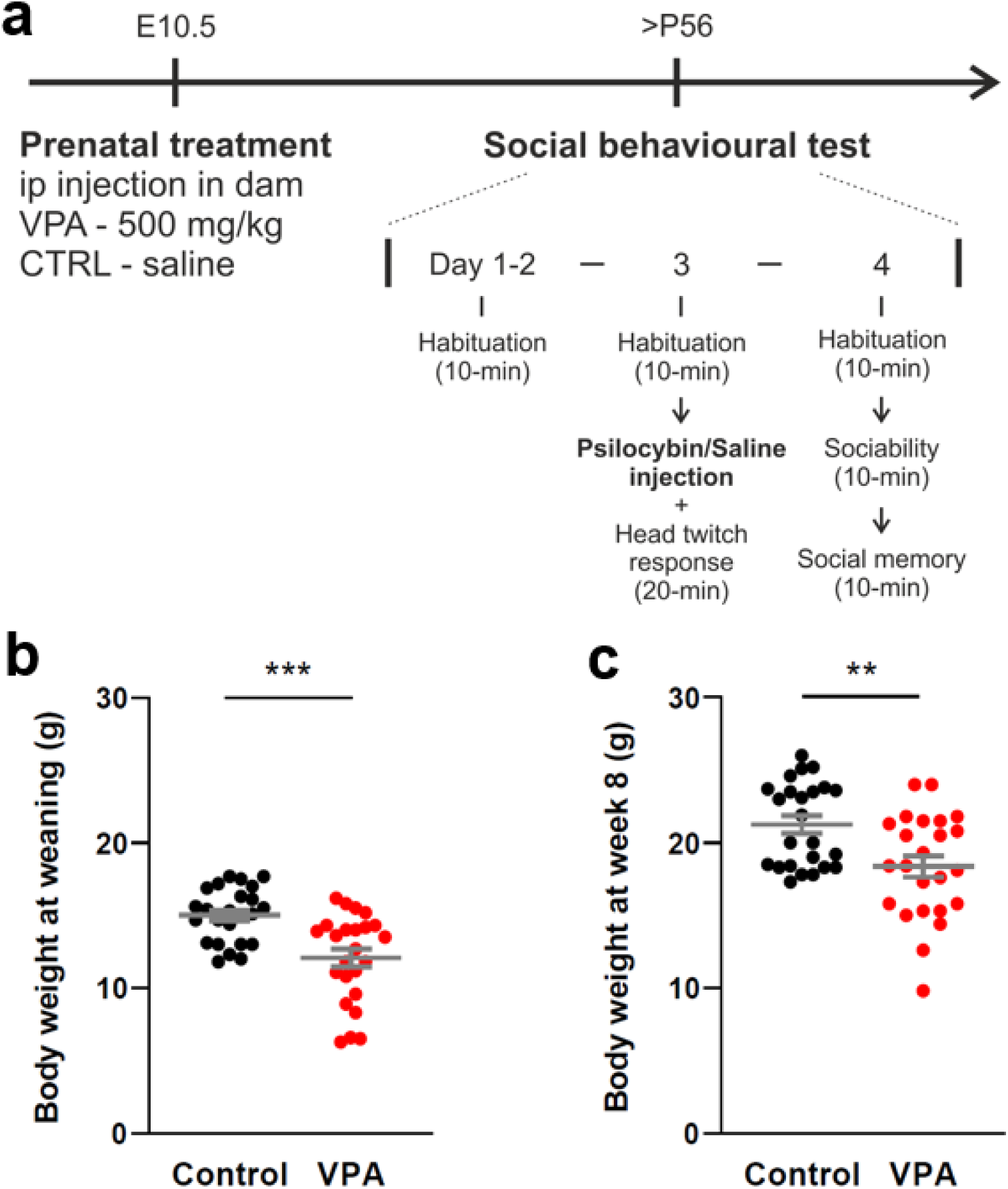
Experimental timeline and VPA model. **a**, Experimental timeline. Pregnant mothers were intraperitoneally injected with saline or 500 mg/kg VPA at E10.5. 8-week old offspring were pre-habituated to the three-chamber arena for three consecutive days. After pre-habituation on day 3, saline or psilocybin was intraperitoneally injected, and the head twitch response (HTR) was counted. On day 4, animals were tested for sociability and social memory using the three-chamber test. **b, c**, Mice from VPA group had lower body weight than the Control group at weaning age (**b**, *t* = 4.033, *P* < 0.001, n = 24/group) and at 8-week old (**c**, *U* = 163.5, *P* = 0.009, n = 24/group). Mean ± SEM. ** *P* < 0.01, *** *P* < 0.001.

**Extended Data Fig. 2:**
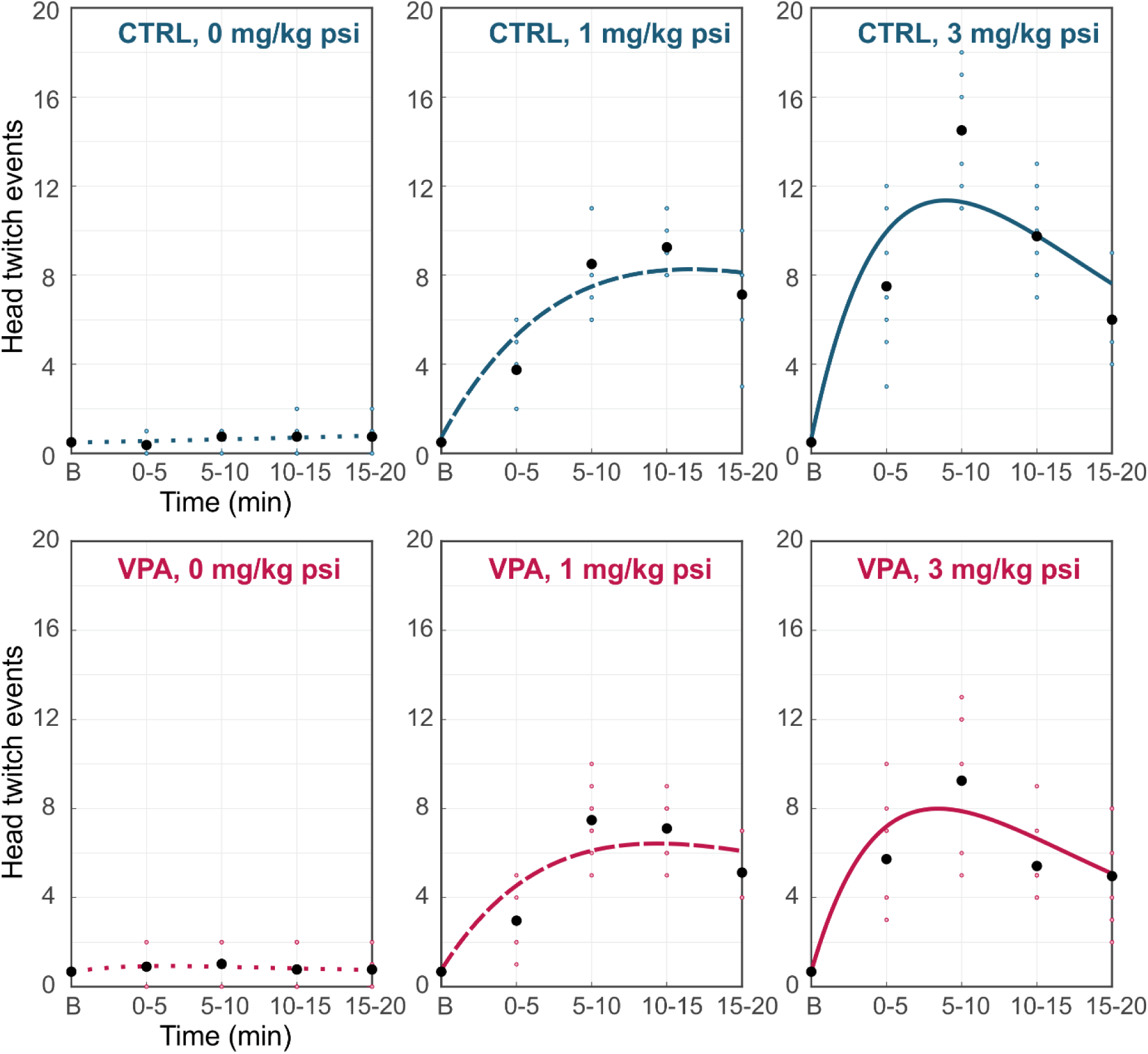
Fitted time course of head twitch events above baseline (B) in 5 min time bins for individual groups. Head twitch events observed in 5-min bins over a total duration of 20-min was fitted with a single dose pharmacokinetics equation. Individual datapoints (blue/pink dots for Control/VPA groups respectively) and group mean values (black dots) are shown. N = 8/group.

## Acknowledgements

We thank Roxanne Wood, Bonnie Glen, and Chantelle Day for their assistance with animal husbandry and care, Imperial Hackspace for their assistance with the behavioural set-up construction, and all members of the Knöpfel lab for discussions. This work was funded by a collaboration grant from COMPASS Pathways, and funding from EU H2020 MSCA 813986 Syn2Psy ITN.

## Author contributions

I.M-G., M.S-R., and C.S. performed the experiments and analysed the data. I.M-G., T.K., T.W., and C.S. conceived the project and wrote the paper. A.Soula, A. Selimbeyoglu and S.H. contributed to the study design.

## Data and code availability

The data and MATLAB scripts that support the findings of this study are available upon reasonable request from the corresponding author.

