## Supplementary Table 1 for "Psilocybin rescues sociability deficits in an animal model of autism"

| | | Comparison group | Test used | N | Descriptive stats | | | P value | | | Degrees of freedom |
|---|---|---|---|---|---|---|---|---|---|---|---|
| | | | | | Group | Mean/Median | SEM | | | | |
| **Physical appearance** | | | | | | | | | | | |
| Extended Fig. 1 | Body weight at weaning | Control vs VPA | Unpaired t-test | 24 | CTRL | 15.01 | 0.38 | t = 4.033 | 0.0002 | SIGNIFICANT | 46 |
| | | | | 24 | VPA | 12.09 | 0.618 | | | | |
| Extended Fig. 1 | Body weight at testing (8wks) | Control vs VPA | Mann-Whitney U | 24 | CTRL | 20.95 | IQR: 18.33-23.68 | U = 163.5 | 0.0095 | SIGNIFICANT | |
| | | | | 24 | VPA | 18.4 | IQR:15.43 - 21.45 | | | | |
| **Head twitch response** | | | | | | | | | | | |
| Fig. 1 | HTR | Pre-treatment: control, VPA x Treatment: saline, psilocybin 1 mg/kg, psilocybin 3 mg/kg | Two-way ANOVA | 48 | Pre-treatment x Treatment | | | F = 4.895 | 0.0123 | SIGNIFICANT | 2,42 |
| | | | Bonferroni's post hoc test | 16 | CTRL_sal vs CTRL_psi1 | | | | < 0.001 | SIGNIFICANT | |
| | | | Bonferroni's post hoc test | 16 | CTRL_sal vs CTRL_psi3 | | | | < 0.001 | SIGNIFICANT | |
| | | | Bonferroni's post hoc test | 16 | VPA_sal vs VPA_psi1 | | | | < 0.001 | SIGNIFICANT | |
| | | | Bonferroni's post hoc test | 16 | VPA_sal vs VPA_psi3 | | | | < 0.001 | SIGNIFICANT | |
| | | | Bonferroni's post hoc test | 16 | CTRL_psi1 vs VPA_psi1 | | | | 0.5179 | NON SIGNIFICANT | |
| | | | Bonferroni's post hoc test | 16 | CTRL_psi3 vs VPA_psi3 | | | | 0.0024 | SIGNIFICANT | |
| | | pre-treatment: control, VPA x Treatment: saline, psilocybin 1 mg/kg, psilocybin 3 mg/kg x Gender: males, females | Three-way ANOVA | 48 | Pre-treatment x Treatment x Gender | | | F = 0.03 | 0.969 | NON SIGNIFICANT | 2,36 |
| **Sociability test** | | | | | | | | | | | |
| Fig. 2a | Total duration of nose-to-nose interactions | Control_saline vs VPA_saline | Unpaired t-test | 7 | CTRL_sal | 26.26 | 3.283 | t = 3.282 | 0.005 | SIGNIFICANT | 15 |
| | | | | 10 | VPA_sal | 12.79 | 2.556 | | | | |
| | | VPA_saline vs VPA_psilocybin 1 mg/kg | Unpaired t-test | 10 | VPA_sal | 12.79 | 2.556 | t = 2.928 | 0.009 | SIGNIFICANT | 18 |
| | | | | 10 | VPA_psi1 | 23.3 | 2.521 | | | | |
| | | VPA_saline vs VPA_psilocybin 3 mg/kg | Unpaired t-test | 10 | VPA_sal | 12.79 | 2.556 | t = 1.071 | 0.2999 | NON SIGNIFICANT | 16 |
| | | | | 8 | VPA_psi3 | 16.61 | 2.38 | | | | |
| | | Control_saline vs Control_psilocybin 1 mg/kg | Unpaired t-test | 7 | CTRL_sal | 26.26 | 3.283 | t = 0.6569 | 0.5227 | NON SIGNIFICANT | 13 |
| | | | | 8 | CTRL_psi1 | 23.61 | 2.457 | | | | |
| | | Control_saline vs Control_psilocybin 3 mg/kg | Unpaired t-test | 7 | CTRL_sal | 26.26 | 3.283 | t =1.799 | 0.0953 | NON SIGNIFICANT | 13 |
| | | | | 8 | CTRL_psi3 | 19.1 | 2.379 | | | | |
| | | Pre-treatment: control, VPA x Treatment: saline, psilocybin 1 mg/kg, psilocybin 3 mg/kg | Two-way ANOVA | 51 | Pre-treatment x Treatment | | | F = 3.633 | 0.0345 | SIGNIFICANT | 2,45 |
| Fig. 2b | Average length of nose-to-nose interactions | Control_saline vs VPA_saline | Unpaired t-test | 7 | CTRL_sal | 1.721 | 0.31 | t = 2.952 | 0.0099 | SIGNIFICANT | 15 |
| | | | | 10 | VPA_sal | 0.903 | 0.088 | | | | |
| | | VPA_saline vs VPA_psilocybin 1 mg/kg | Unpaired t-test | 10 | VPA_sal | 0.903 | 0.088 | t = 3.307 | 0.0039 | SIGNIFICANT | 18 |
| | | | | 10 | VPA_psi1 | 1.388 | 0.117 | | | | |
| | | VPA_saline vs VPA_psilocybin 3 mg/kg | Unpaired t-test | 10 | VPA_sal | 0.903 | 0.088 | t = 1.326 | 0.2033 | NON SIGNIFICANT | 16 |
| | | | | 8 | VPA_psi3 | 1.11 | 0.136 | | | | |
| | | Control_saline vs Control_psilocybin 1 mg/kg | Unpaired t-test | 7 | CTRL_sal | 1.721 | 0.31 | t = 1.698 | 0.115 | NON SIGNIFICANT | 12 |
| | | | | 8 | CTRL_psi1 | 1.164 | 0.105 | | | | |
| | | Control_saline vs Control_psilocybin 3 mg/kg | Unpaired t-test | 7 | CTRL_sal | 1.721 | 0.31 | t = 1.911 | 0.078 | NON SIGNIFICANT | 13 |
| | | | | 8 | CTRL_psi3 | 1.131 | 0.101 | | | | |
| | | Pre-treatment: control, VPA x Treatment: saline, psilocybin 1 mg/kg, psilocybin 3 mg/kg | Two-way ANOVA | 50 | Pre-treatment x Treatment | | | F = 6.657 | 0.003 | SIGNIFICANT | 2,44 |
| **Social memory test** | | | | | | | | | | | |
| Fig. 2c | Social novelty index | Pre-treatment: control, VPA x Treatment: saline, psilocybin 1 mg/kg, psilocybin 3 mg/kg | Two-way ANOVA | 43 | Pre-treatment x Treatment | | | F = 6.077 | 0.0052 | SIGNIFICANT | 2,37 |
| | | Pre-treatment: control, VPA x Treatment: saline, psilocybin 1 mg/kg, psilocybin 3 mg/kg | Bonferroni's post hoc test | 14 | CTRL_sal vs CTRL_psi3 | | | | 0.0276 | SIGNIFICANT | |
| | | Pre-treatment: control, VPA x Treatment: saline, psilocybin 1 mg/kg, psilocybin 3 mg/kg | Bonferroni's post hoc test | 16 | CTRL_psi1 vs CTRL_psi3 | | | | 0.047 | SIGNIFICANT | |
| | | Pre-treatment: control, VPA x Treatment: saline, psilocybin 1 mg/kg, psilocybin 3 mg/kg | Bonferroni's post hoc test | 15 | CTRL_psi3 vs VPA_psi3 | | | | 0.0057 | SIGNIFICANT | |
